# QPot: An R Package for Stochastic Differential Equation Quasi-Potential Analysis

**DOI:** 10.1101/029777

**Authors:** Christopher M. Moore, Christopher R. Stieha, Ben C. Nolting, Maria K. Cameron, Karen C. Abbott

## Abstract

**QPot** (pronounced 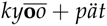) is an R package for analyzing two-dimensional systems of stochastic differential equations. It provides users with a wide range of tools to simulate, analyze, and visualize the dynamics of these systems. One of **QPot**’s key features is the computation of the quasi-potential, an important tool for studying stochastic systems. Quasi-potentials are particularly useful for comparing the relative stabilities of equilibria in systems with alternative stable states. This paper describes **QPot**’s primary functions, and explains how quasi-potentials can yield insights about the dynamics of stochastic systems. Three worked examples guide users through the application of **QPot**’s functions.

## Introduction

Differential equations are an important modeling tool in virtually every scientific discipline. Most differential equation models are deterministic, meaning that they provide a set of rules for how variables change over time, and no randomness comes into play. Reality, of course, is filled with random events (i.e., noise or stochasticity). Unfortunately, many of the analytic techniques developed for deterministic ordinary differential equations are insufficient to study stochastic systems, where phenomena like noise-induced transitions between alternative stable states and metastability can occur. For systems subject to stochasticity, the quasi-potential is a tool that yields information about properties such as the expected time to escape a basin of attraction, the expected frequency of transitions between basins, and the stationary probability distribution. **QPot** (abbreviation of **Q**uasi-**P**otential) is an R package that allows users to calculate quasi-potentials, and this paper is a guided tutorial of its application.

Key functions

| Function | Main arguments | Description |
| --- | --- | --- |
| TSTraj() | Deterministic skeleton, $\sigma$, $T$, $\Delta t$ | Creates a realization (time series) of the stochastic differential equations. |
| TSPlot() | TSTraj() output | Plots a realization of the stochastic differential equations, with an optional histogram side-plot. Plots can additionally be two-dimensional, which show realizations in $(X, Y)$-space. |
| TSDensity() | TSTraj() output | Creates a density plot of a trajectory in $(X, Y)$-space in one or two dimensions. |
| QPotential() | Deterministic skeleton, stable equilibria, bounds, mesh (number of divisions along each axis) | Creates a matrix corresponding to a discretized version of the local quasi-potential function for each equilibrium. |
| QPGlobal() | Local quasi-potential matrices, unstable equilibria | Creates a global quasi-potential surface. |
| QPInterp() | Global quasi-potential, $(x, y)$-coordinates | Evaluates the global quasi-potential at $(x, y)$. |
| QPContour() | Global quasi-potential | Creates a contour plot of the quasi-potential. |
| VecDecomAll() | Global quasi-potential, deterministic skeleton, bounds | Creates three vector fields: the deterministic skeleton, the negative gradient of the quasi-potential, and the remainder vector field. To find each field individually, the functions `VecDecomVec()`, `VecDecomGrad()`, or `VecDecomRem()` can be used. |
| VecDecomPlot() | Deterministic skeleton, gradient, or remainder field | Creates a vector field plot for the vector, gradient, or remainder field. |

## Adding stochasticity to deterministic models

Consider a differential equation model of the form

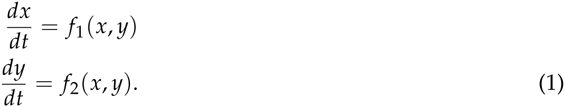

In many cases, state variables are subject to continual random perturbations, which are commonly modeled as white noise processes. To incorporate these random influences, the original system of deterministic differential equations can be transformed into a system of stochastic differential equations:

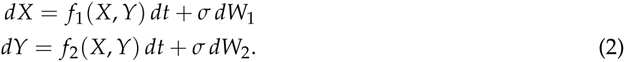

*X* and *Y* are now stochastic processes (a change emphasized through the use of capitalization); this means that, at every time *t*, *X*(*t*) and *Y*(*t*) are random variables, as opposed to real numbers. *σ* ≥ 0 is a parameter specifying the noise intensity, and W_1_ and W_2_ are Wiener processes. A Wiener process is a special type of continuous-time stochastic process whose changes over non-overlapping time intervals, Δ*t*_1_ and Δ*t*_2_, are independent Gaussian random variables with mean zero and variances 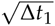 and 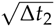, respectively. The differential notation in equations (2) is a formal way of representing a set of stochastic integral equations, which must be used because realizations of Wiener processes are not differentiable (to be precise, with probability one, a realization of a Wiener process will be almost nowhere differentiable). The functions *f*_1_ and *f*_2_ are called the deterministic skeleton. The deterministic skeleton can be viewed as a vector field that determines the dynamics of trajectories in the absence of stochastic effects. We will forgo a complete overview of stochastic differential equations here; interested readers are encouraged to seek out texts like Allen (2007) and Iacus (2009). We note, however, that throughout this paper we use the Itô formulation of stochastic differential equations.

## The quasi-potential

Consider system (2), with deterministic skeleton (1). If there exists a function *V* (*x, y*) such that 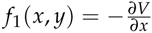 and 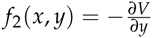, then system (1) is called a gradient system and *V* (*x, y*) is called the system’s potential function. The dynamics of a gradient system can be visualized by considering the (*x*, *y*)-coordinates of a ball rolling on a surface specified by *z* = *V*(*x, y*). Gravity causes the ball to roll downhill, and stable equilibria correspond to the bottoms of the surface’s valleys. *V* (*x, y*) is a Lyapunov function for the system, which means that if (*x*(*t*), *y*(*t*)) is a solution to the system of equations (1), then 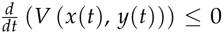, and the only places that 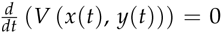 are at equilibria. This means that the ball’s elevation will monotonically decrease, and will only be constant if the ball is at an equilibrium. The basin of attraction of a stable equilibrium **e**\* of system (1) is the set of points that lie on solutions that asymptotically approach **e**\*.

The potential function is useful for understanding the stochastic system (2). As in the deterministic case, the dynamics of the stochastic system can be represented by a ball rolling on the surface *z* = *V* (*x, y*); in the stochastic system, however, the ball experiences random perturbations due to the noise terms in system (2). In systems with multiple stable equilibria, these random perturbations can cause a trajectory to move between different basins of attraction. The depth of the potential (that is, the difference in *V* at the equilibrium and the lowest point on the boundary of its basin of attraction), is a useful measure of the stability of the equilibrium (see Nolting and Abbott, Accepted). The deeper the potential, the less likely it will be for stochastic perturbations to cause an escape from the basin of attraction. This relationship between the potential and the expected time to escape from a basin of attraction can be made precise (formulae in the appendices of Nolting and Abbott, Accepted). Similarly, the potential function is directly related to the expected frequency of transitions between different basins, and to the stationary probability distribution of system (2).

Unfortunately, gradient systems are very special, and a generic system of the form (1) will almost certainly not be a gradient system. That is, there will be no function *V* (*x, y*) that satisfies 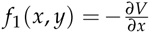 and 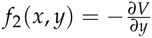. Fortunately, quasi-potential functions generalize the concept of a potential function for use in non-gradient systems. The quasi-potential, Φ(*x*, *y*), of a non-gradient system will possess many of the important properties that potential functions have in gradient systems. The surface *z* = Φ(*x*, *y*) describes the system’s dynamics, and the depth of the surface is a highly useful stability metric. Analogous to the potential described above, Φ(*x*, *y*) is directly related to the the stationary probability distribution, the expected frequency of transitions between basins of attraction, and the expected time required to escape each basin.

In both this paper and in **QPot**, the function that we refer to as the quasi-potential is 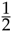 times the quasi-potential as defined by Freidlin and Wentzell (Freidlin and Wentzell, 2012). This choice is made so that the quasi-potential will agree with the potential in gradient systems.

**QPot** is an R package that contains tools for calculating and analyzing quasi-potentials. The following three examples show how to use the tools in this package. The first example is a simple consumer-resource model from ecology. This example is explained in detail, starting with the analysis of the deterministic skeleton, proceeding with simulation of the stochastic system, and finally demonstrating the calculation, analysis, and interpretation of the quasi-potential. The second and third examples are covered in less detail, but illustrate some special system behaviors. Systems with limit cycles, like example 2, require a slightly different procedure than systems that only have point attractors. Extra care must be taken constructing global quasi-potentials for exotic systems, like example 3. For more information about quasi-potentials, see Cameron (2012), Nolting and Abbott (Accepted), and the references therein.

## Example 1: A consumer-resource model with alternative stable states

Consider the stochastic version (*sensu* (2)) of a standard consumer-resource model of plankton (*X*) and their consumers (*Y*) (Collie and Spencer, 1994; Steele and Henderson, 1981):

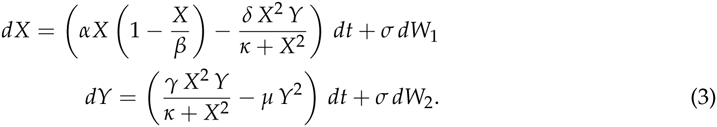

The model is formulated with a Type III functional response, meaning that the highest per-capita consumption rate of plankton occurs at intermediate plankton densities. *α* is the plankton’s maximum population growth rate, *β* is the plankton carrying capacity, *δ* is the maximal feeding rate of the consumers, *γ* is the conversion rate of plankton to consumer, and *μ* is the consumer mortality rate. We will analyze this example with a set of parameter values that yield two stables states: *α =* 1.54, *β =* 10.14, *γ =* 0.476, *δ* = 1, *κ =* 1, and *μ =* 0.112509.

### Step 1: Analyzing the deterministic skeleton

There are preexisting tools in R for analyzing the deterministic skeleton of system (3), which will be described briefly in this subsection. The first step is to find the equilibria for the system and determine their stability with linear stability analysis. Equilibria can be found using the package **rootSolve** (Soetaert and Herman, 2008). In example 1, the equilibria are **e***_u_*_1_ = (0,0), **e***_s_*_1_ = (1.4049, 2.8081), **e***_u_*_2_ = (4.2008, 4.0039), **e***_u_*_2_ = (4.9040, 4.0619), and **e***_u_*_3_ = (10.14, 0). The package **deSolve** (Soetaert et al., 2010) can find the eigenvalues of the linearized system at an equilibrium, which determines the asymptotic stability of the system. **e***_u_*_1_ is an unstable source and **e***_u_*_2_ and **e***_u_*_3_ are saddles. The eigenvalues corresponding to **e***_s_*_1_ are −0.047 ± 0.548 *i* and the eigenvalues corresponding to **e***_u_*_2_ are −0.377 and −0.093. Hence **e***_s_*_1_ is a stable spiral point and **e***_s_*_2_ is a stable node. To ease transition from packages such as **deSolve** to our package **QPot**, we include the wrapper function `Model2String()`, which takes a function containing equations and a list of parameters and their values, and returns the equations in a string that is usable by **QPot**.

The package **phaseR** (Grayling, 2014) generates a stream plot of the deterministic skeleton of the system of equations (3) (Figure 1). Further, **deSolve** can be used to find solutions corresponding to particular initial conditions of the deterministic skeleton of system (3). During the analysis of the deterministic skeleton of a system, it is important to note several things. The first is the range of *x* and *y* values over which relevant dynamics occur. In example 1, transitions between the stable equilibria are a primary point of interest, so one might wish to focus on a region like the one displayed in Figure 1, even though this region excludes **e***_u_*_3_. The ranges of the variables will determine the window sizes and ranges used later in the quasi-potential calculations. Second, it is important to note if there are any limit cycles. If there are, it will be necessary to identify a point on the limit cycle. This can be accomplished by calculating a long-time solution of the system of ODEs to obtain a trajectory that settles down on the limit cycle (see example 2). Finally, it is important to note regions of phase space that correspond to unbounded solutions. As explained in subsequent sections, it is worth examining system behavior in negative phase space, even in cases where negative quantities lack physical meaning.

**Figure 1:**
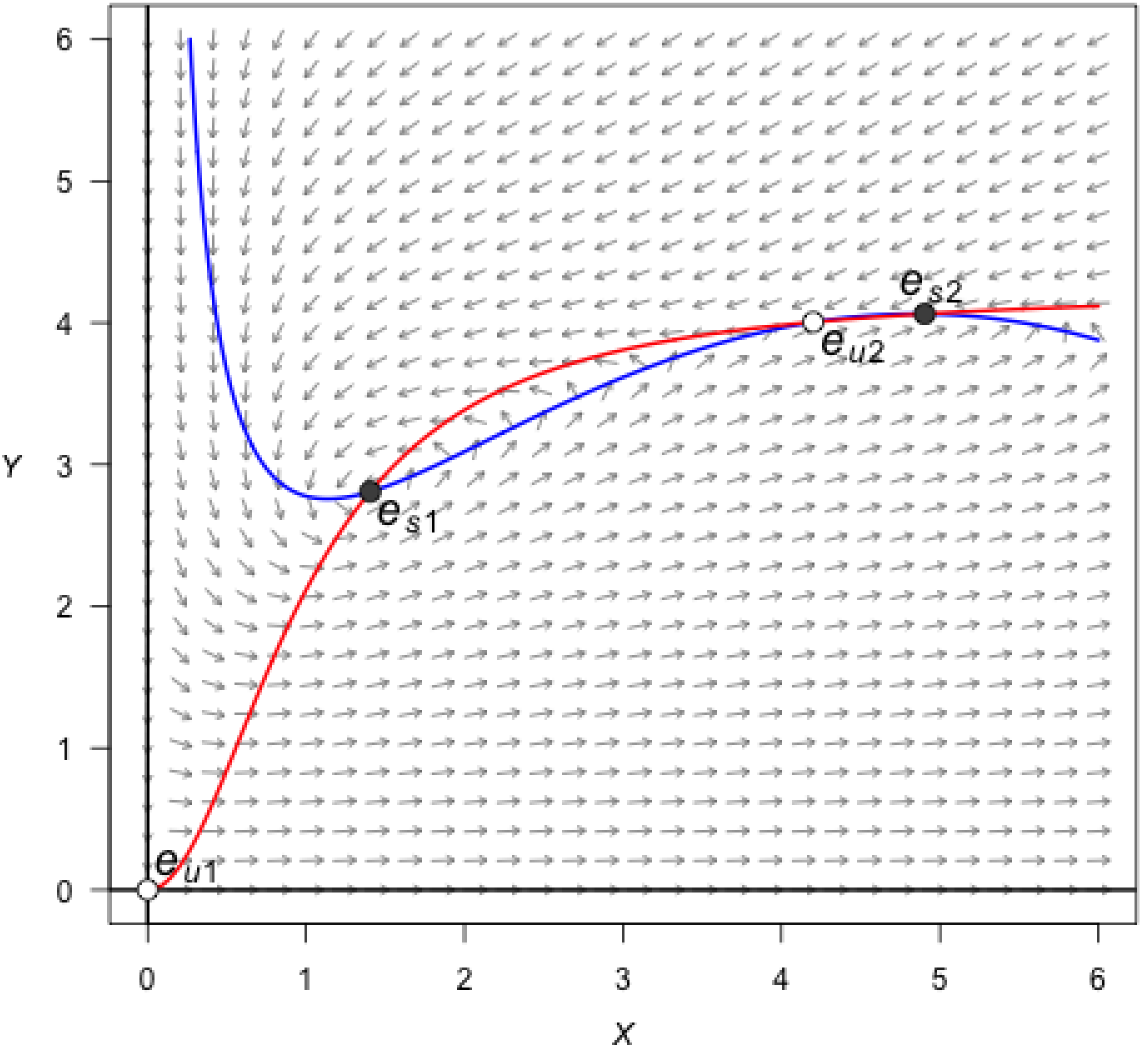
A stream plot of the deterministic skeleton of system (3). The blue line is an *x*-nullcline (where 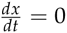) and the red line is a *y*-nullcline (where 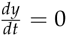). Open circles are unstable equilibria and filled circles are stable equilibria. Made using the package **phaseR.**

### Step 2: Stochastic simulation

For a specified level of noise intensity, *σ*, one can obtain a realization of system (3). To do this, `TSTraj()` in **QPot** implements the Euler-Maruyama method. All other code/function references hereafter are found in **QPot**, unless specified otherwise. To generate a realization, the following arguments are required: the right-hand side of the deterministic skeleton for both equations, the initial conditions (*x*_0_, *y*_0_), the parameter values, the step-size Δ*t*, and the total time length *T*.

~~~
var.eqn.x <- “(alpha*x)*(1-(x/beta)) - ((delta*(x^^^2)*y)/(kappa+(x^^^2)))”
var.eqn.y <- “((gamma*(x^^^2)*y)/(kappa+(x^^^2))) - mu*(y^^^2)”
model.state <- c(x = 1, y = 2)
model.parms <- c(alpha = 1.54, beta = 10.14, delta = 1, gamma = 0.476, kappa =1, mu = 0.112509)
model.sigma <- 0.05
model.time <- 1000 # we used 12500 in the figures
model.deltat <- 0.025
ts.ex1 <- TSTraj(y0 = model.state, time = model.time, deltat = model.deltat, x.rhs = var.eqn.x, y.rhs = var.eqn.y, parms = model.parms, sigma = model.sigma)
~~~

Figure 2 shows a realization for *σ =* 0.05, Δ*t* = 0.025, *T =* 1.25 × 10^4^, and initial condition (*x*_0_, *y*_0_) = (1, 2). The argument `dim = 1` produces a time series plot with optional histogram side-plot. The `dim = 2` produces a plot of a realization in (*x, y*)-space. If the system is ergodic, a very long realization will approximate the steady-state probability distribution. Motivated by this, a probability density function can be approximated from a long realization using the `TSDensity()` function (e.g., Figure 3).

**Figure 2:**
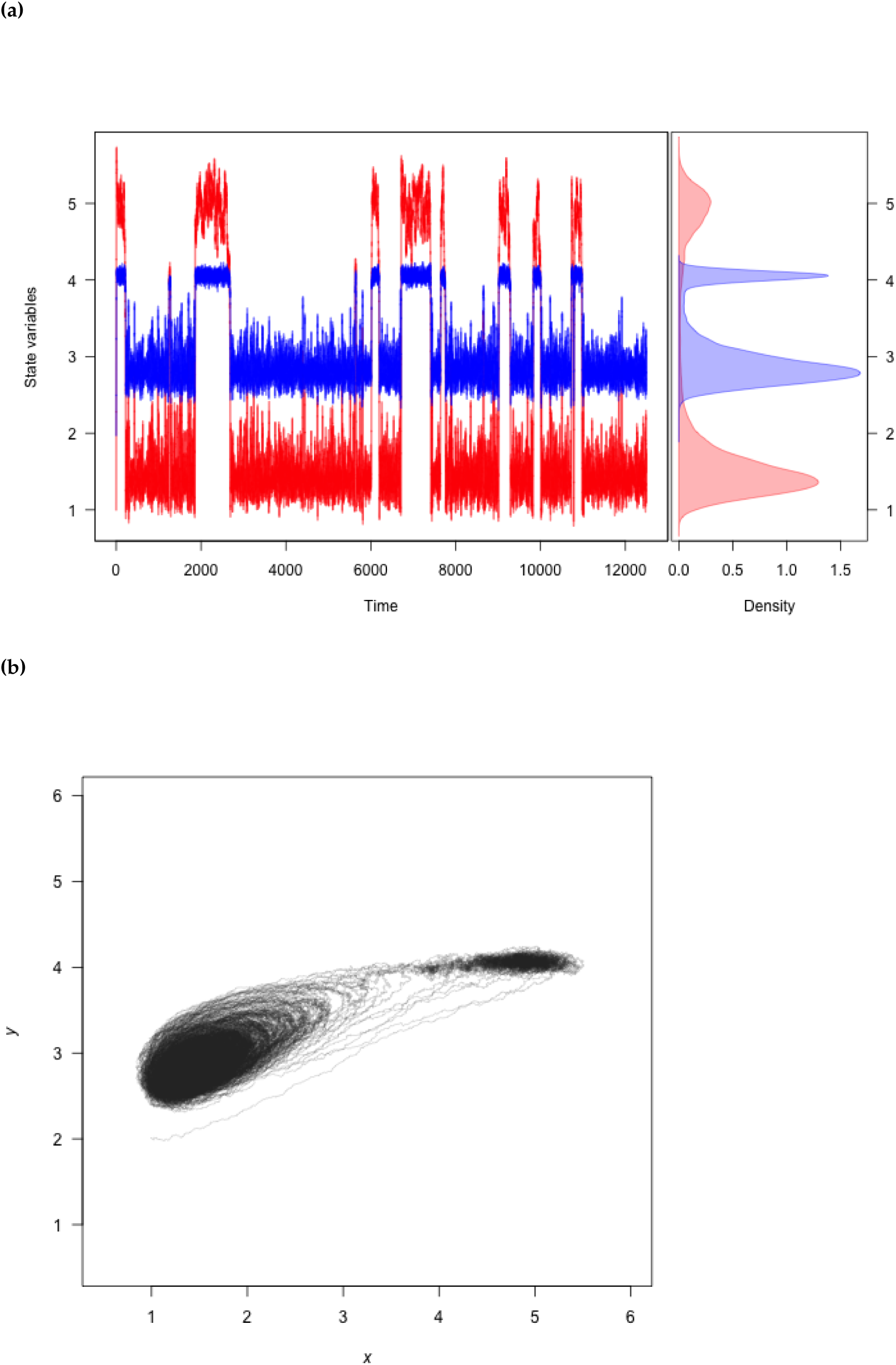
**(a)** A realization of system (3) created using `TSPlot()`, with *x* in blue and *y* in red. The left side of (a) shows the time series. The right side of (a), which is enabled with the default dens = T, shows a histogram of the *x* and *y* values over the entire realization. **(b)** The realization plotted in (*x, y*)-space with `dim = 2`.

**Figure 3:**
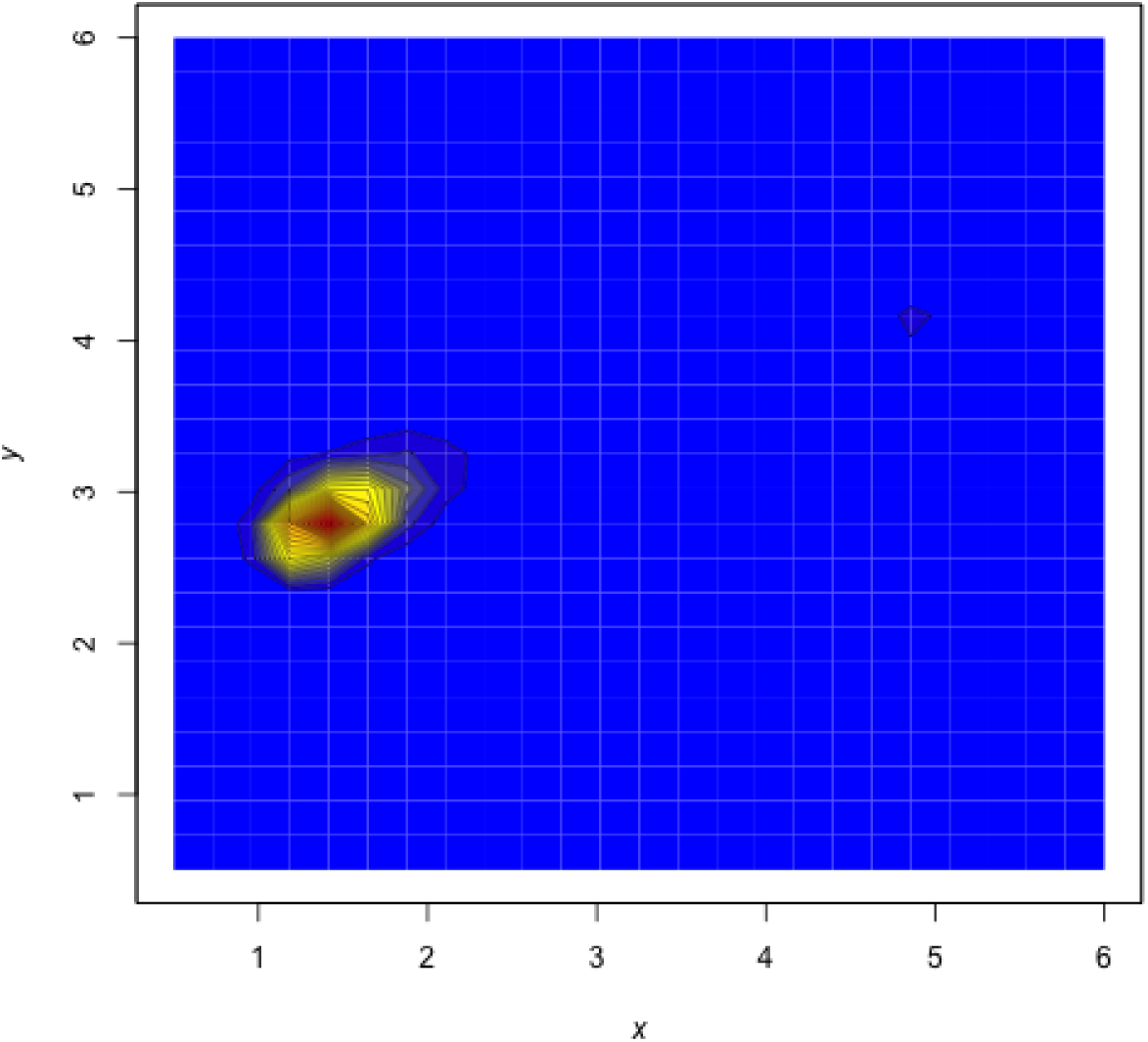
A density plot obtained from a realization of system (3). Red corresponds to high density, and blue to low density. Plotted using the function `TSDensity() with dim = 2`.

~~~
TSPlot(ts.ex1, deltat = model.deltat)                       # Figure 2a
TSPlot(ts.ex1, deltat = model.deltat, dim =2)          # Figure 2a
TSDensity(ts.ex1, dim =1)                      # like Figure 2a histogram
TSDensity(ts.ex1, dim =2)                                 # Figure 3
~~~

Bounds can be placed on the state variables in all of the functions described in this subsection. For example, it might be desirable to set 0 as the minimum size of a biological population, because negative population densities are not physically meaningful. A lower bound can be imposed on the functions described in this subsection with the argument `lower`. bound in the function `TRTraj()`. Similarly, it might be desirable to set an upper bound for realizations, and hence prevent runaway trajectories (unbounded population densities are also not physically meaningful). An upper bound can be imposed on the functions described in this subsection with the argument upper.bound.

### Step 3: Local quasi-potential calculation

The next step is to compute a local quasi-potential for each attractor. Because **QPot** deals with two-dimensional systems, “attractor” will be used synonymously with “stable equilibrium” “or stable limit cycle”. A limit cycle will be considered in example 2. For now, suppose that the only attractors are stable equilibrium points, **e***_s_*_i_, *i =* 1, &, *n*. In the example above, *n =* 2. For each stable equilibrium **e***_si_*, we will compute a local quasi-potential Φ*_i_*(*x, y*).

In order to understand the local quasi-potential, it is useful consider the analogy of a particle traveling according to system (2). In the context of example 1, the coordinates of the particle correspond to population densities, and the particle’s path corresponds to how those population densities change over time. The deterministic skeleton of (2) can be visualized as a force field influencing the particle’s trajectory. Suppose that the particle moves along a path from a stable equilibrium **e***_si_* to a point (*x, y*). If this path does not coincide with a solution of the deterministic skeleton, then the stochastic terms must be doing some “work” to move the particle along the path. The more work that is required, the less likely it is for the path to be a realization of system (2). Φ*_i_*(*x*, *y*) is the amount of work required to traverse the easiest path from **e***_si_* to (*x, y*). Note that Φ*_i_*(*x*, *y*) is non-negative, and it is zero at **e***_s_*_i_.

In the basin of attraction for **e***_si_*, Φ*_i_*(*x*, *y*) has many properties analogous to the potential function for gradient systems. Key among these properties is that the quasi-potential is non-increasing along deterministic trajectories. This means that the quasi-potential can be interpreted as a type of energy surface, and the rolling ball metaphor is still valid. The difference is that, in non-gradient systems, there is an additional component to the vector field that causes trajectories to circulate around level sets of the energy surface. This is discussed in more detail in Step 6, below.

**QPot** calculates quasi-potentials using an adjustment developed by Cameron (2012) to the ordered upwind algorithm (Sethian and Vladimirsky, 2001, 2003). The idea behind the algorithm is to calculate Φ*_i_*(*x*, *y*) in ascending order, starting with the known point **e***_si_*. The result is an expanding area where the solution is known.

Calculating Φ*_i_*(*x*, *y*) with the function `QPotential()` requires a text string of the equations and parameter values, the stable equilibrium points, the computation domain, and the mesh size. For (3), this first means inputting the equations:

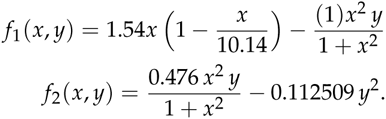

In R:

~~~
equation.x = “1.54*x*(1.0-(x/10.14))-(y*x*x)/(1.0+x*x)”
equation.y = “((0.476*x*x*y)/(1+x*x))-0.112590*y*y”
~~~

The coordinates of the points **e***_si_*, which were determined in Step 1, are **e***_s_*_1_ = (1.4049, 2.8081) and **e***_s_*_2_ = (4.9040, 4.0619).

~~~
eq1.x = 1.40491
eq1.y = 2.80808
eq2.x = 4.9040
eq2.y = 4.06187
~~~

Next, the boundaries of the computational domain need to be entered. This domain will be denoted by [L*x*_1_, L*x*_2_] × [L*y*_1_, *Ly*_2_]. The ordered-upwind method terminates when the solved area encounters a boundary of this domain. Thus, it is important to choose boundaries carefully. For example, if **e***_si_* lies on one of the coordinate axes, one should not use that axis as a boundary because the algorithm will immediately terminate. Instead, one should add padding space. This is important even if the padding space corresponds to physically unrealistic values (e.g., negative population densities). For this example, a good choice of boundaries is: L*x*_1_ = L*y*_1_ = −0.5, and *Lx*_2_ *= Ly*_2_ *=* 20. This choice of domain was obtained by examining stream plots of the deterministic skeleton and density plots of stochastic realizations (Figures 1–3). The domain contains all of the deterministic skeleton equilibria, and it encompasses a large area around the regions of phase space visited by stochastic trajectories (Figures 1–3). Note that a small padding space was added to the left and bottom sides of the domain, so that the coordinate axes are not the domain boundaries.

~~~
bounds.x = c(-0.5, 20.0)
bounds.y = c(-0.5, 20.0)
~~~

In some cases, it may be desirable to treat boundaries differently in the upwind algorithm. This is addressed below in the section “Boundary behavior”.

Finally, the mesh size for the discretization of the domain needs to be specified. Let *N_x_* be the number of grid points in the *x*-direction and *N_y_* be the number of grid points in the *y*-direction. Note that the horizontal distance between mesh points is 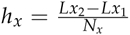, and the vertical distance between mesh points is 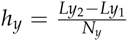. Mesh points are considered adjacent if their Euclidean distance is less than or equal to 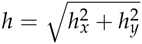. This means that diagonal mesh points are considered adjacent. In this example, a good choice is *N_x_ = Ny =* 4100. This means that *h_x_ = hy =* 0.005, and *h* ≈ 0.00707. In general, the best choice of mesh size will be a compromise between resolution and computational time. The mesh size must be fine enough to precisely track how information moves outward along characteristics from the initial point. Too fine of a mesh size can lead to very long computational times, though. The way that computation time scales with grid size depends on the system under consideration (see below for computation time for this example), because the algorithm ends when it reaches a boundary, which could occur before the algorithm has exhaustively searched the entire mesh area.

~~~
step.number.x = 1000 # we used 4100 in the figures
step.number.y = 1000 # we used 4100 in the figures
~~~

The “anisotropy ratio” is another adjustable parameter for the algorithm, defined by k.x and k.y in `QPotential()`. For more on this, see Cameron (2012). For now, we suggest using the defaults *K_x_ =* 20 and *K_y_ =* 20.

The R interface implements the `QPotential()` algorithm using C code. By default `QPotential()` outputs a matrix that contains the quasi-potentials to the R session. The time required to compute the quasi-potential will depend on the size of the region and the fineness of the mesh. This example with *K_x_ = K_y_ =* 20 and *N_x_ = N_y_ =* 4100 has approximately 1.7 × 10^7^ grid points, which leads to run times of approximately 2.25 min (2.5 GHz Intel Core i5 processor and 8 GB 1600 MHz DDR3 memory). When one reaches around 5 × 10^8^, computational time can be several hours. Setting the argument `save.to.R` to `TRUE` outputs the matrix into the R session, and setting the argument `save.to.HD` to `TRUE` saves the matrix to the hard drive as the file `filename` in the current working directory. For *N_x_ = N_y_ =* 4100, the saved file occupies 185 MB.

~~~
eq1.local <- QPotential(x.rhs = equation.x, x.start = eq1.x, x.bound = bounds.x, x.num.steps = step.number.x, y.rhs = equation.y, y.start = eq1.y, y.bound = bounds.y, y.num.steps = step.number.y)
Step 3 should be repeated until local quasi-potentials Φ*_i_*(*x*, *y*) have been obtained for each **e***_si_*. In example 1, this means calculating Φ_1_(*x*, *y*) corresponding to **e***_s_*_1_ and Φ_2_(*x*, *y*) corresponding to **e***_s_*_2_.
eq2.local <- QPotential(x.rhs = equation.x, x.start = eq2.x, x.bound = bounds.x, x.num.steps = step.number.x, y.rhs = equation.y, y.start = eq2.y, y.bound = bounds.y, y.num.steps = step.number.y)
~~~

Each local quasi-potential Φ*_i_*(*x*, *y*) is stored in R as a large matrix. The entries in this matrix are the values of Φ*_i_*, at each mesh point. To define the function on the entire domain (i.e., to allow it to be evaluated at arbitrary points in the domain, not just the discrete mesh points), bilinear interpolation is used. The values of Φ(*x*, *y*) can be extracted using the function `QPInterp()`. Inputs to `QPInterp()` include the (*x, y*) coordinates of interest, the (*x, y*) domain boundaries, and the `QPotential()` output (i.e., the matrix with rows corresponding to *x*-values and columns corresponding to *y*-values). `QPInterp()` can be used for any of the local quasi-potential or the global quasi-potential surfaces (see the next subsection).

### Step 4: Global quasi-potential calculation

Recall that Φ*_i_*(*x*, *y*) is the amount of “work” required to travel from **e***_si_* to (*x, y*). This information is useful for considering dynamics in the basin of attraction of **e***_si_*. In many cases, however, it is desirable to define a global quasi-potential that describes the system’s dynamics over multiple basins of attraction. If a gradient system has multiple stable states, the potential function provides an energy surface description that is globally valid. We seek an analogous global function for non-gradient systems. Achieving this requires “pasting” local quasi-potentials into a single global quasi-potential. If the system has only two attractors, one can define a global quasi-potential, though, it might be nontrivial, see example 3 ahead. In systems with three or more attractors such a task might not be possible (Freidlin and Wentzell, 2012). For a wide variety of systems, however, a relatively simple algorithm can accomplish the pasting (Graham and Tél, 1986; Roy and Nauman, 1995). In most cases, the algorithm amounts to translating the local quasi-potentials up or down so that they agree at the saddle points that separate the basins of attraction. In example 1, **e***_u_*_1_ lies on the boundary of the basins of attraction for **e***_s_*_1_ and **e***_s_*_2_. Creating a global quasi-potential requires matching Φ_1_ and Φ_2_ at **e***_u_*_2_. Φ_1_(**e***_u_*_2_) = 0.007056 and Φ_2_(**e***_u_*_2_) = 0.00092975. If one defines

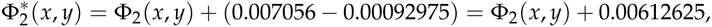

then Φ_1_ and 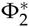 match at **e***_u_*_2_. Finally, define

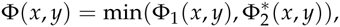

which is the global quasi-potential. For systems with more than two stable equilibria, this process is generalized to match local quasi-potentials at appropriate saddles. QPot automates this procedure. A fuller description of the underlying algorithm is explained in example 3, which requires a more nuanced understanding of the pasting procedure.

~~~
ex1.global <- QPGlobal(local.surfaces = list(eq1.local, eq2.local), unstable.eq.x = c(0, 4.2008), unstable.eq.y = c(0, 4.0039), x.bound = bounds.x, y.bound = bounds.y)
~~~

This function `QPGlobal` calculates the global quasi-potential by automatically pasting together the local quasi-potentials. This function requires the input of all the discretized local quasi-potentials, and the coordinates of all of the unstable equilibria. The output is a discretized version of the global quasi-potential. The length of time required for this computation will depend on the total number of mesh points; for the parameters used in example 1, it takes a couple of minutes. As with the local quasi-potentials, the values of Φ(*x*, *y*) can be extracted using the function `QPInterp()`.

### Step 5: Global quasi-potential visualization

To visualize the global quasi-potential, one can simply take the global quasi-potential matrix from `QPGlobal` and use it to create a contour plot using `QPContour()` (Figure 4).

**Figure 4:**
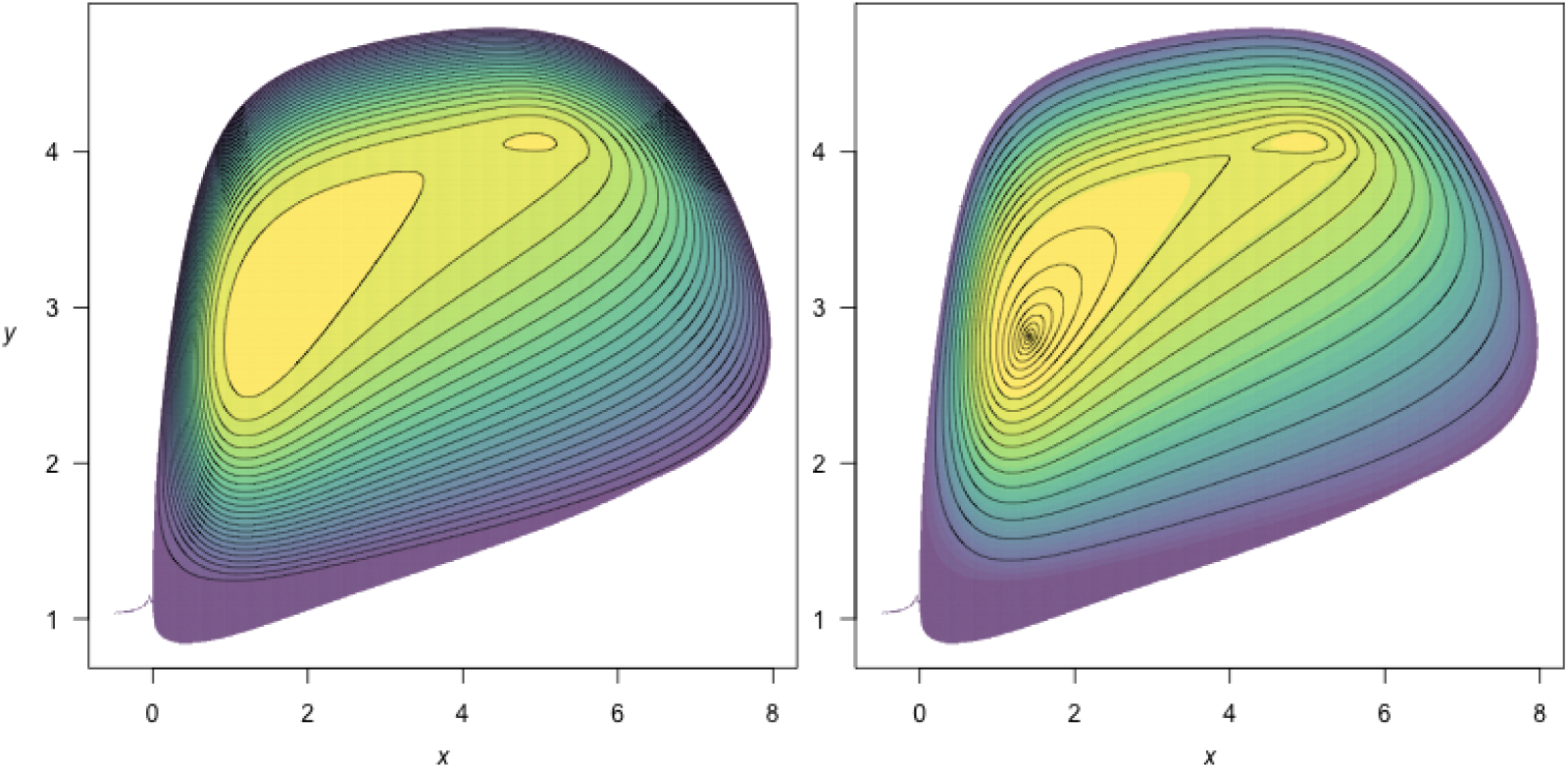
A contour plot of the of the quasi-potential of system (3). Yellow corresponds to low values of the quasi-potential, and purple to high values. `c.parm` in `QPContour()`, can be used to condense the contour lines at the bottom of the basins for better resolution. The default creates evenly spaced contour lines (left; `c.parm = 1`). On the right, contour lines are condensed (`c.parm = 5`).

**Figure 5:**
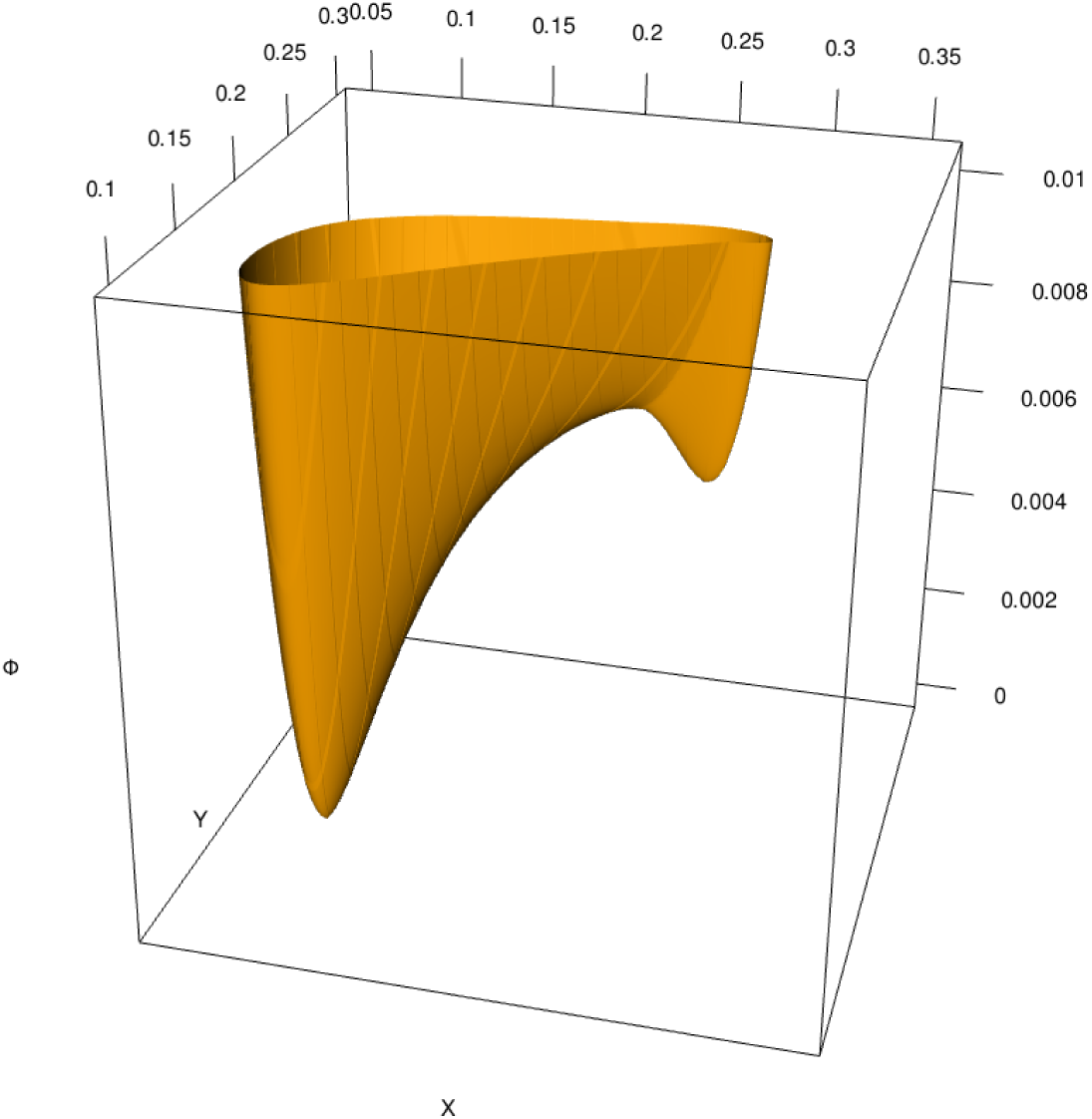
A 3D plot of the of the quasi-potential of system (3) using `persp3d()` in package **rg1**. 3D plotting can further help users visualize the quasi-potential surfaces.

~~~
QPContour(surface = ex1.global, dens = c(1000, 1000), x.bound = bounds.x, y.bound = bounds.y, c.parm = 5) # right side of Figure~\ref{fig:ex1qp}
~~~

`QPContour()` is based on the `.filled.contour()` function from the base package **graphics**. In most cases, the mesh sizes used for the quasi-potential calculation will be much finer than what is required for useful visualization. The argument dens within `QPContour()` reduces the points used in the graphics generation. Although it might seem wasteful to perform the original calculations at a mesh size that is finer than the final visualization, this is not so. Choosing the mesh size in the original calculations to be very fine reduces the propagation of errors in the ordered upwind algorithm, and hence leads to a more accurate numerical solution.

An additional option allows users to specify contour levels. R’s default for the `contour()` function creates contour lines that are equally spaced over the range of values specified by the user. In some cases, however, it is desirable to use a non-linear spacing for the contours. For example, equally-spaced contours will not capture the topography at the bottom of a basin if the changes in height are much smaller than other regions in the plot. Simply increasing the number of equally-spaced contour lines does not solve this problem, because steep areas of the plot become completely saturated with lines. `QPContour()` has a function for non-linear contour spacing that condenses contour lines at the bottoms of basins. Specifically, for *n* contour lines, this function generates a list of contour levels, 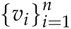, specified by:

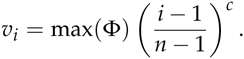

*c =* 1 yields evenly-spaced contours. As *c* increases, the contour lines become more concentrated near basin bottoms. Figure 4 shows equal contour lines (left panel) and contour lines that are concentrated at the bottom of the basin (right panel, c.parm = 5).

Finally, creating a 3D plot can be very useful for visualizing the features of more complex surfaces. This is especially helpful when considering the physical metaphor of a ball rolling on a surface specified by a quasi-potential (Nolting and Abbott, Accepted). R has several packages for 3D plotting, including static plotting with the base function persp() and with the package **plot3D** (Soetaert, 2013). Interactive plotting is provided by **rg1** (Adler et al., 2015). To create an interactive 3D plot for example 1 using **rg1**, use the code: `persp3d(x = ex1.global,col = “orange”)`. Figure shows a 3D plot of example 1 that clearly illustrates the differences between the two local basins. Users can also export the matrix of quasi-potential values and create 3D plots in other programs.

### Step 6: Vector field decomposition

Recall that the deterministic skeleton (1) can be visualized as a vector field, as shown in Figure 1. In gradient systems, this vector field is completely determined by the potential function, *V*(*x, y*). The name “gradient system” refers to the fact that the vector field is the negative of the potential function’s gradient,

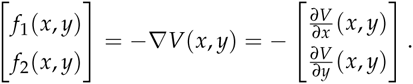

In non-gradient systems, the vector field can no longer be represented solely in terms of the gradient of Φ(*x, y*). Instead, there is a remainder component of the vector field, 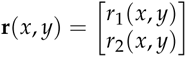. The vector field can be decomposed into two terms:

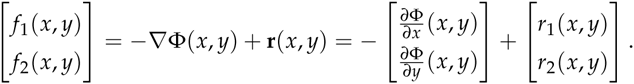

The remainder vector field is orthogonal to the gradient of the quasi-potential everywhere. That is, for every (*x*, *y*) in the domain,

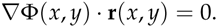

An explanation of this property can be found in Nolting and Abbott (Accepted).

The remainder vector field can be interpreted as a force that causes trajectories to circulate around level sets of the quasi-potential. **QPot** enables users to perform this decomposition. The function `VecDecomAll()` calculates the vector field decomposition, and outputs three vector fields: the original deterministic skeleton, **f** (*x, y*); the gradient vector field, −∇Φ (*x, y*); and the remainder vector field, **r**(*x, y*). Each of these three vector fields can be output alone using `VecDecomDS()`, `VecDecomGrad()`, or `VecDecomRem()`. These vector fields can be visualized using the function `VecDecomPlot()`. Code to create the vector fields from `VecDecomAll()` is displayed below; code for generating individual vector fields can be found in `help()` for `VecDecomDS()`, `VecDecomGrad()`, or `VecDecomRem()`. The gradient and remainder vector fields are shown in the left and right columns of figure 6, respectively, with proportional vectors (top row) and equal-length vectors (bottom row). Three arguments within `VecDecomPlot()` are important to creating comprehensible plots: dens, tail.length, and head.length. dens specifies the number of arrows in the plot window along the *x* and *y* axes. The argument `tail.length` scales the length of arrow tails. The argument `head.length` scales the length of arrow heads. The function `arrows()` makes up the base of `VecDecomPlot()`, and arguments can be passed to it, as well as to plot. The code below produces all three vector fields from the multi-dimensional array returned by `VecDecomAll()`:

**Figure 6:**
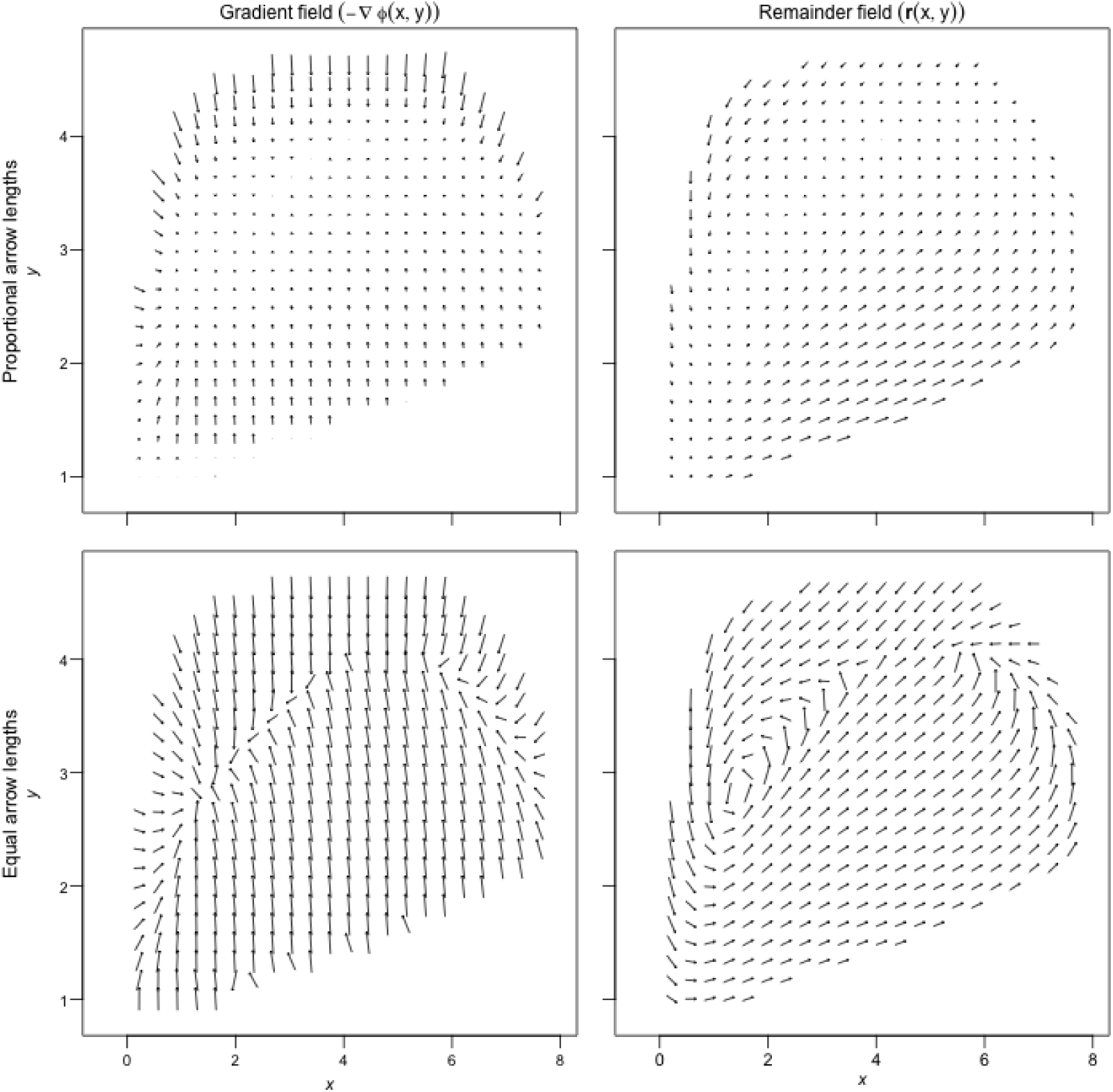
The gradient (left column) and remainder (right column) fields, plotted with `arrow.type = “proportional”` (top row) and `arrow.type = “equal”` (bottom row) arrow lengths using `VecDecomPlot()` for system (3).

~~~
# Calculate all three vector fields
VDAll <- VecDecomAll(surface = ex1.global, x.rhs = equation.x, y.rhs = equation.y, x.bound = bounds.x, y.bound = bounds.y)
# Plot the deterministic skeleton vector field
VecDecomPlot(field = list(VDAll[,,1], VDAll[,,2]), dens = c(25, 25),
x.bound = bounds.x, y.bound = bounds.y, x.lim = c(0, 11), y.lim = c(0, 6), arrow.type = “equal”, tail.length = 0.25, head.length = 0.025)
# Plot the gradient vector field
VecDecomPlot(field = list(VDAll[,,3], VDAll[,,4]), dens = c(25, 25),
x.bound = bounds.x, y.bound = bounds.y, arrow.type = “proportional”, tail.length = 0.25, head.length = 0.025)
# Plot the remainder vector field
VecDecomPlot(field = list(VDAll[,,5], VDAll[,,6]), dens = c(25, 25),
x.bound = bounds.x, y.bound = bounds.y, arrow.type = “proportional”, tail.length = 0.35, head.length = 0.025)
~~~

## Example 2: A model with a limit cycle

Consider the following model:

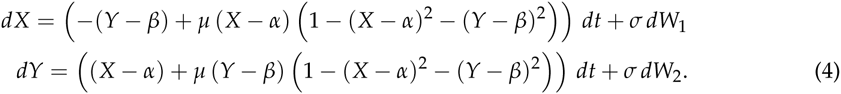

We will analyze this example with *μ =* 0.2, *α =* 4, and *β =* 5.

### Step 1: Analyzing the deterministic skeleton

The deterministic skeleton of this system has one equilibrium, **e**_0_ = (4, 5), which is an unstable spiral point. Figure 7 shows a stream plot of the deterministic skeleton of system (4). A particular solution of the deterministic skeleton of system (4) can be found using **rootSolve** and **deSolve**. The stream plot and a few particular solutions suggest that there is a stable limit cycle. To calculate the limit cycle, once can find a particular solution over a long time interval (e.g., Figure 7 has three trajectories run for *T =* 100). The solution will eventually converge to the limit cycle. One can drop the early part of the trajectory until only the closed loop of the limit cycle remains. There are more elegant ways to numerically find a periodic orbit (even when those orbits are unstable). For more information on these methods, see Chua and Parker (1989). In this example, the limit cycle is shown by the thick black line in Figure 7. For calculation of the quasi-potential, it is sufficient to input a single point that lies on the limit cycle. For this example, one such point is **z** = (4.15611, 5.98774).

**Figure 7:**
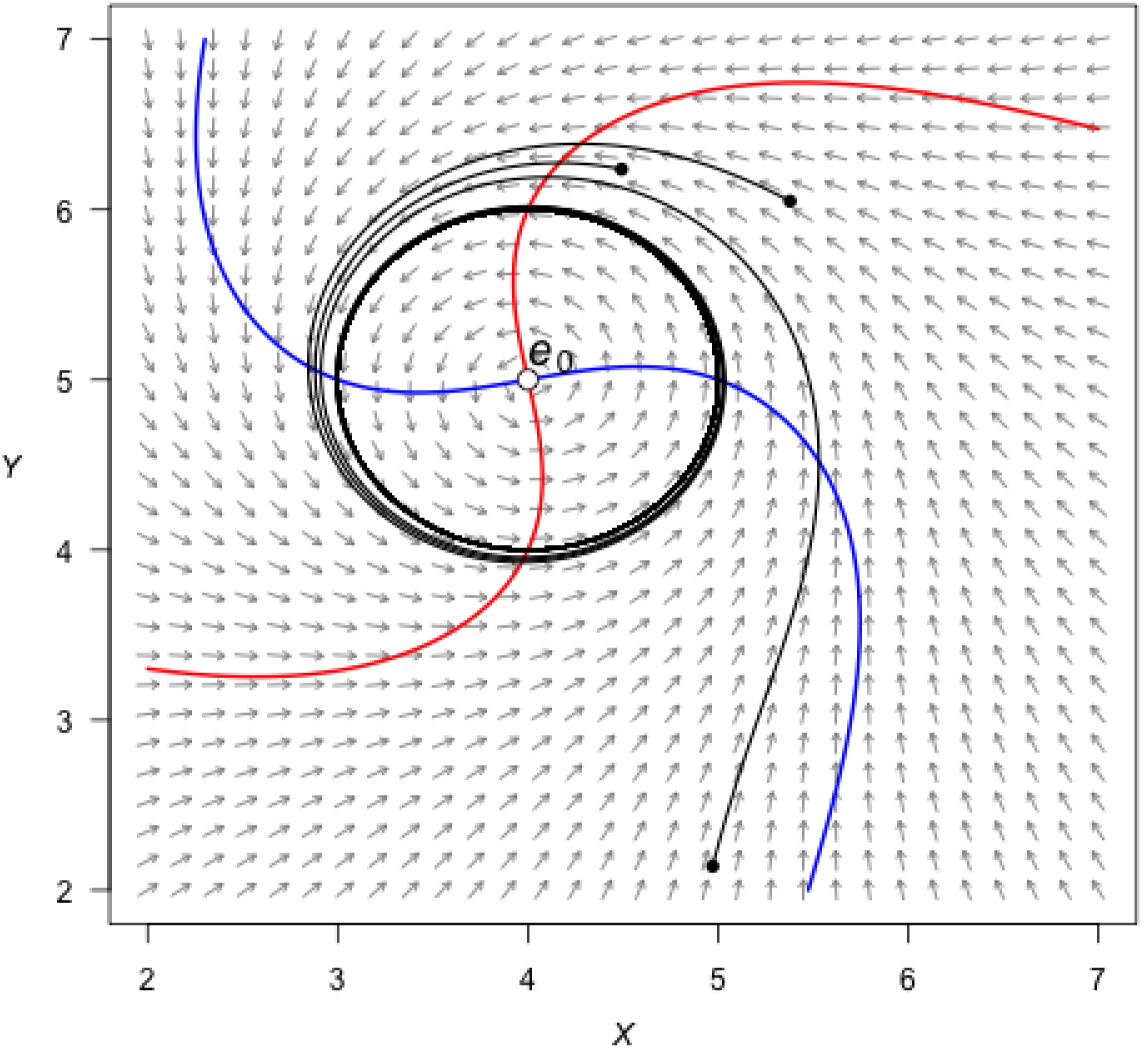
A stream plot of the deterministic skeleton of system (4). The blue line is an *x*-nullcline (where 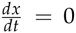) and the red line is a *y*-nullcline (where 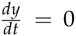). The open circle is an unstable equilibrium. Particular solutions are shown as black lines, with filled circles as initial conditions. Made using the package **phaseR**.

### Step 2: Stochastic simulation

Figure 8 shows a time series for a realization of (4) with *σ =* 0.1, Δ*t =* 5 × 10^−3^, *T* = 250 and initial condition (*x*_0_, *y*_0_) = (3, 3). Figure 9 shows a density plot of a realization with the same parameters, except *T* = 2.5 × 10^3^.

**Figure 8:**
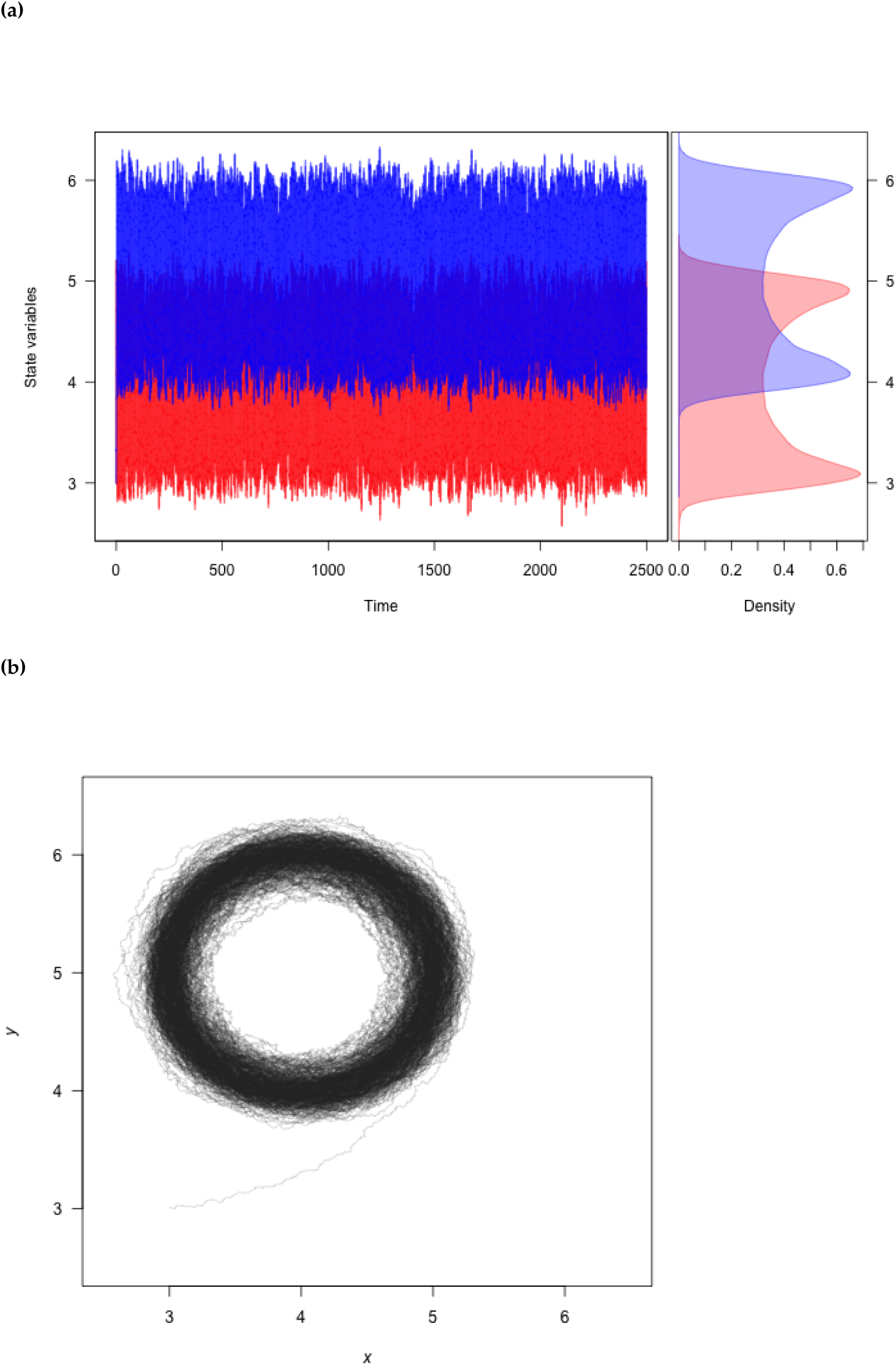
**(a)** A realization of system (4) created using `TSPlot()`, with x in blue and y in red. The left side of (a) shows the time series. The right side of (a), which is enabled with the default dens = TRUE, shows a histogram of the *x* and *y* values over the entire realization. **(b)** The realization plotted in (*x, y*)-space (`dim = 2` in the function `TSPlot()`).

**Figure 9:**
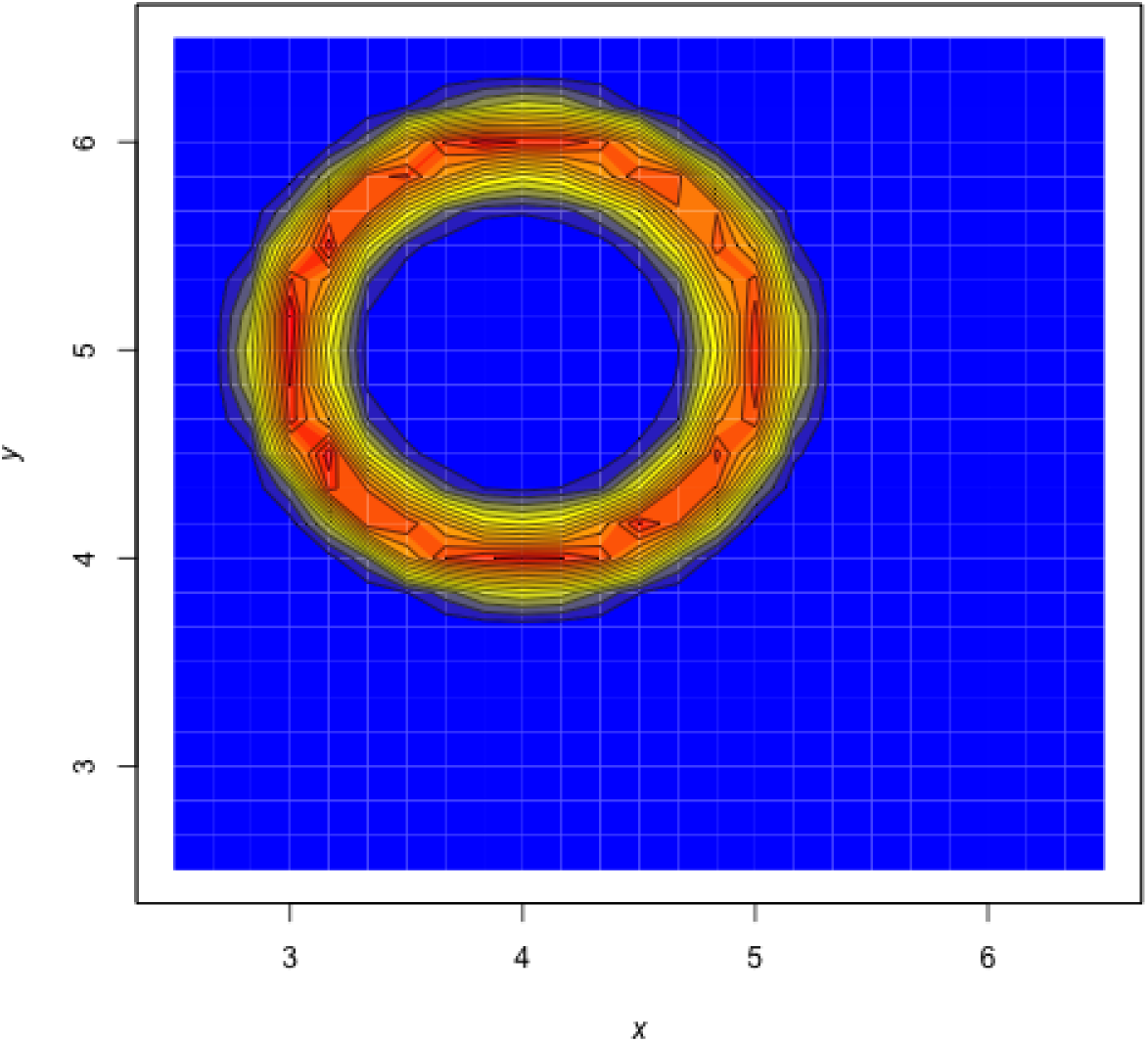
A density plot obtained from a realization of system (4) using `TSDensity()` with `dim = 2`. Red corresponds to high density, and blue to low density.

~~~
var.eqn.x <- “-(y-beta) + mu*(x-alpha)*(1-(x-alpha)^^^2-(y-beta)^^^2)”
var.eqn.y <- “(x-alpha) + mu*(y-beta)*(1-(x-alpha)^^^2-(y-beta)^^^2)”
model.state <- c(x = 3, y = 3)
model.parms <- c(alpha = 4, beta = 5, mu = 0.2)
model.sigma <-0.1
model.time <- 1000 # we used 2500 in the figures
model.deltat <- 0.005
ts.ex2 <- TSTraj(y0 = model.state, time = model.time, deltat = model.deltat, x.rhs = var.eqn.x, y.rhs = var.eqn.y, parms = model.parms, sigma = model.sigma)
TSPlot(ts.ex2, deltat = model.deltat)                       # Figure 8a
TSPlot(ts.ex2, deltat = model.deltat, dim = 2, line.alpha = 25)                # Figure 8b
TSDensity(ts.ex2, dim =1)            # Histogram
TSDensity(ts.ex2, dim =2)            # Figure 9
~~~

### Step 3: Local quasi-potential calculation

In this example, there are no stable equilibrium points. There is one stable limit cycle, and this can be used to obtain a local quasi-potential. Using **z** as the initial point for the ordered-upwind algorithm and L*x*_1_ = −0.5, *Ly*_1_ = −0.5, L*x*_2_ = 7.5, *Ly*_2_ = 7.5, *Nx* = 4000, and *Ny* = 4000, one obtains a local quasi-potential, Φ_z_(*x, y*). This generates the local quasi-potential Φ_z_(*x, y*).

~~~
eqn.x <- “-(y-5) + (0.2)*(x-4)*(1-(x-4)^^^2-(y-5)^^^2)”
eqn.y <- “(x-4) + (0.2)*(y-5)*(1-(x-4)^^^2-(y-5)^^^2)”
eq1.qp <- QPotential(x.rhs = eqn.x, x.start = 4.15611, x.bound = c(-0.5, 7.5), x.num.steps = 4000, y.rhs = eqn.y, y.start = 5.98774, y.bound = c(-0.5, 7.5), y.num.steps = 4000)
~~~

### Step 4: Global quasi-potential calculation

There is only one local quasi-potential in this example, so it is the global quasi-potential, Φ(*x, y*) =Φ_z_(*x, y*).

### Step 5: Global quasi-potential visualization

Figure 10 shows a contour plot of the global quasi-potential.

**Figure 10:**
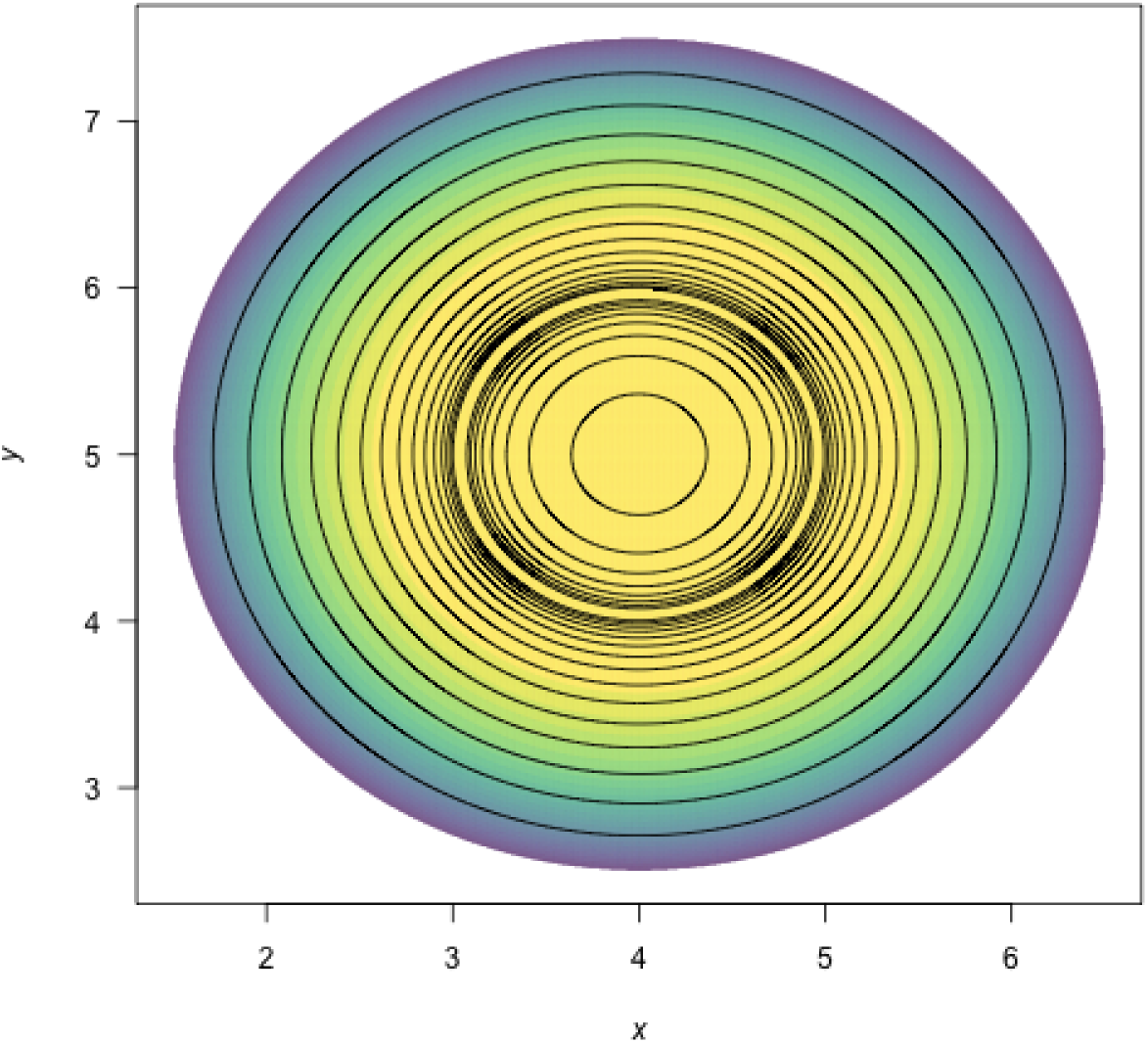
A contour plot of the of the quasi-potential of system (4) using `QPContour()`. Yellow corresponds to low values of the quasi-potential, and purple to high values.

~~~
QPContour(eq1.qp, dens = c(1000, 1000), x.bound = c(-0.5, 7.5), y.bound = c(-0.5, 7.5), c.parm = 10)
~~~

## Example 3: More complicated local quasi-potential pasting

In example 1, the procedure for pasting local quasi-potentials together into global quasi-potential was a simple, two-step process. First, one of the local quasi-potentials was translated so that the two surfaces agreed at the saddle point separating the two basins of attraction. Second, the global quasi-potential was obtained by taking the minimum of the two surfaces at each point. A general algorithm for pasting local quasi-potentials, as explained in Graham and Tél (1986) and Roy and Nauman (1995), is slightly more complicated. This process is automated in QPGlobal, but it is worth understanding the process in order to correctly interpret the outputs.

To understand the full algorithm, consider the following model:

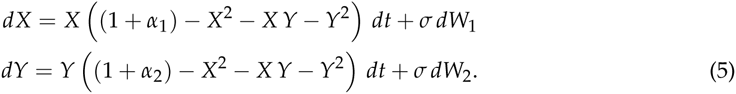

For this analysis, let *α*_1_ = 1.25 and *α*_2_ = 2.

### Step 1: Analyzing the deterministic skeleton

The deterministic skeleton of this system has five equilibria. These are **e***_u_*_1_ = (0,0), **e***_s_*_1_ = (0, −1.73205), **e***_s_*_2_ = (0, 1.73205), **e***_u_*_2_ = (−1.5, 0) and **e***_u_*_3_ = (1.5, 0). The eigenvalue analysis shows that **e***_u_*_1_ is an unstable node, **e***_s_*_1_ and **e***_s_*_2_ are stable nodes, **e***_u_*_2_ and **e***_u_*_3_ are saddles. Figure 11 shows a stream plot of the deterministic skeleton of (5). The basin of attraction for **e***_s_*_1_ is the lower half-plane, and the basin of attraction for **e***_s_*_2_ is the upper half-plane.

**Figure 11:**
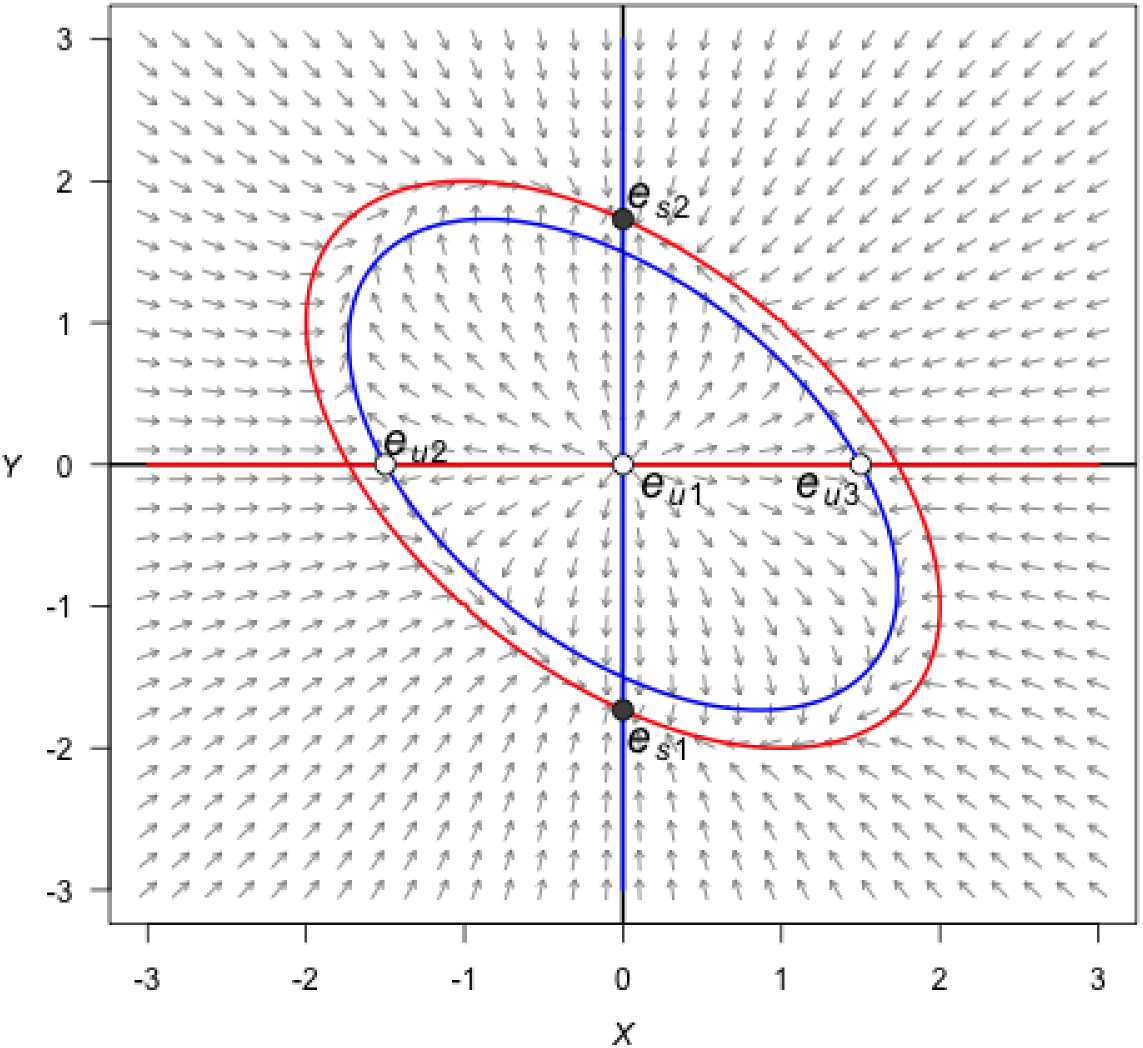
A stream plot of the deterministic skeleton of system (5). The blue line is an *x*-nullcline (where 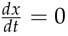) and the red line is a *y*-nullcline (where 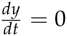). Open circles are stable equilibria and filled circles are unstable equilibria. Made using the package **phaseR**.

### Step 2: Stochastic simulation

Figure 12 shows a time series for a realization of system (5) with *σ =* 0.8, Δ*t* = 0.01, *T* = 5000 and initial condition (*x*_0_, *y*_0_) = (0.5, 0.5). Figure 13 shows a density plot of this realization.

**Figure 12:**
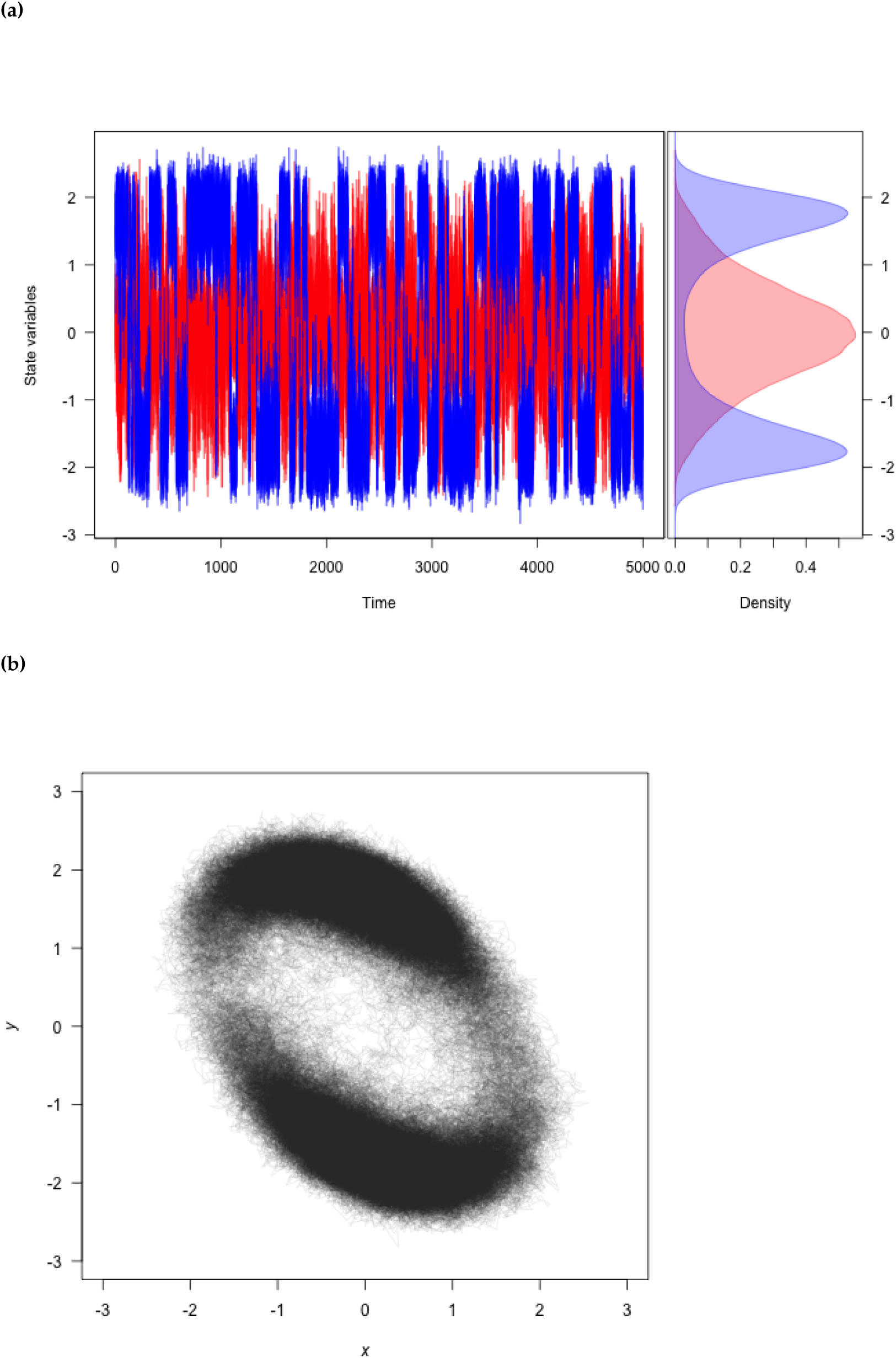
**(a)** A realization of system (5) created using `TSPlot()`, with *x* in blue and *y* in red. The left side in panel (a) shows the time series. The right side in panel (a), which is enabled by default with parameter `dens = T` in the function `TSPlot()`, shows a histogram of the *x* and *y* values over the entire realization. **(b)** The realization plotted in (*x, y*)-space with `TSPlot()` with `dim = 2`.

**Figure 13:**
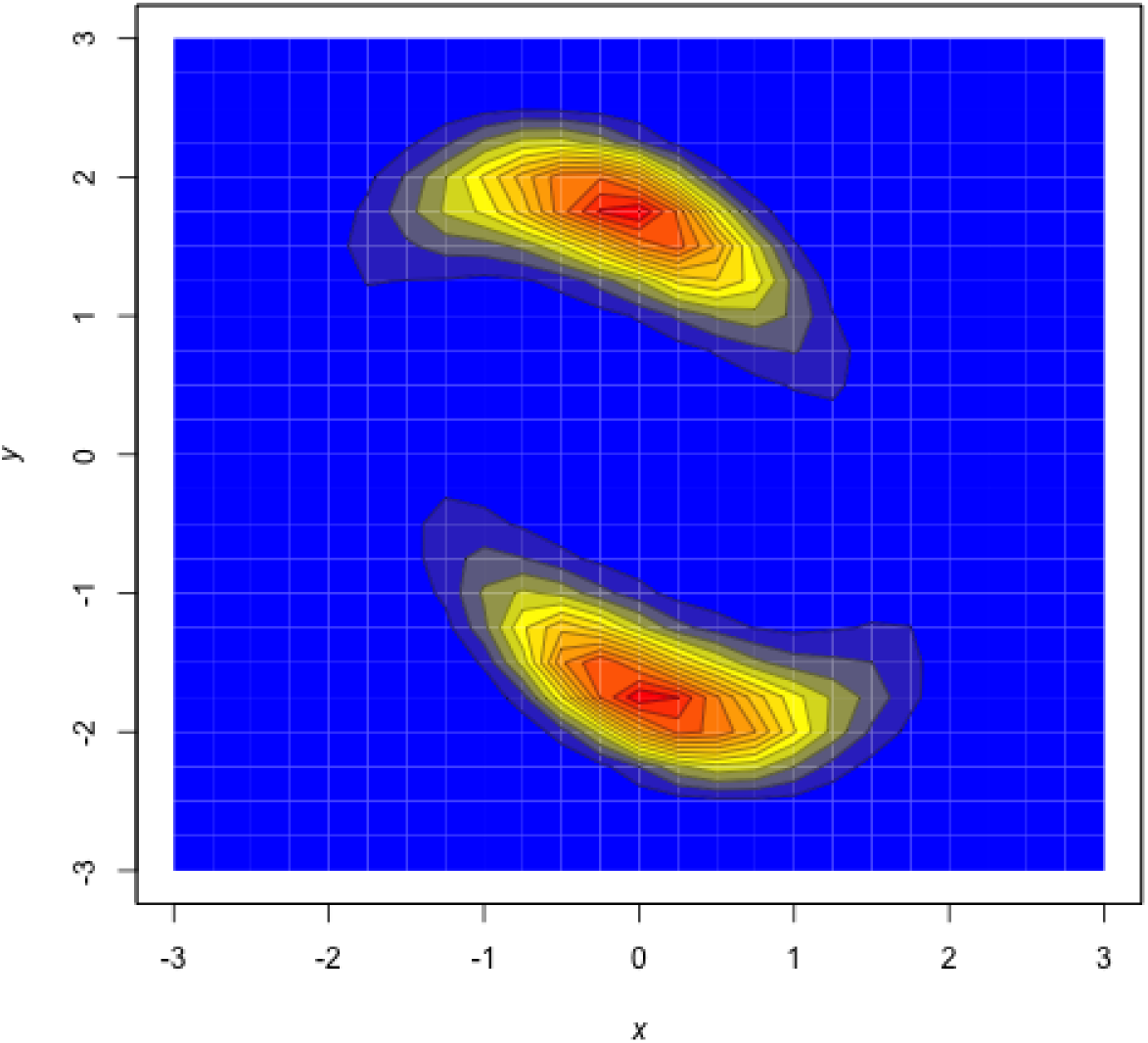
A density plot obtained from a realization of system (5) by using the function `TSDensity()` with `dim = 2, contour.levels = 20`, and `contour.lwd = 0.1`. Red corresponds to high density, and blue to low density.

~~~
var.eqn.x <- “x*((1+alpha1)-x*x-x*y-y*y)”
var.eqn.y <- “y*((1+alpha2)-x*x-x*y-y*y)”
model.state <- c(x = 0.5, y = 0.5)
model.parms <- c(alpha1 = 1.25, alpha2 = 2)
model.sigma <- 0.8
model.time <- 5000
model.deltat <- 0.01
ts.ex3 <- TSTraj(y0 = model.state, time = model.time, deltat = model.deltat, x.rhs = var.eqn.x, y.rhs = var.eqn.y, parms = model.parms, sigma = model.sigma)
TSPlot(ts.ex3, deltat = model.deltat)                                             # figure 12a
TSPlot(ts.ex3, deltat = model.deltat, dim = 2, line.alpha = 25)         # figure 12b
TSDensity(ts.ex3, dim =1)                                             # Histogram of time series
TSDensity(ts.ex3, dim = 2, contour.levels = 20, contour.lwd = 0.1) # figure 13
~~~

### Step 3: Local quasi-potential calculation

Two local quasi-potentials need to be calculated, Φ_1_(*x, y*) corresponding to **e***_s_*_1_, and Φ_2_(*x*, *y*) corresponding to **e***_s_*_2_. In both cases, sensible boundary and mesh choices are L*x*_1_ = −3, L*y*_1_ = −3, *Lx*_2_ = 3, *Ly*_2_ = 3, *Nx* = 6000, and *Ny* = 6000.

~~~
equation.x = “x*((1+1.25)-x*x-x*y-y*y)”
equation.y = “y*((1+2)-x*x-x*y-y*y)”
bounds.x = c(-3, 3)
bounds.y = c(-3, 3)
step.number.x = 6000
step.number.y = 6000
eq1.x = 0
eq1.y = -1.73205
eq2.x = 0
eq2.y = 1.73205
eq1.local <- QPotential(x.rhs = equation.x, x.start = eq1.x, x.bound = bounds.x, x.num.steps = step.number.x, y.rhs = equation.y, y.start = eq1.y, y.bound = bounds.y, y.num.steps = step.number.y)
eq2.local <- QPotential(x.rhs = equation.x, x.start = eq2.x, x.bound = bounds.x, x.num.steps = step.number.x, y.rhs = equation.y, y.start = eq2.y, y.bound = bounds.y, y.num.steps = step.number.y)
~~~

### Step 4: Global quasi-potential

If one were to naively try to match the local quasi-potentials at **e***_u_*_2_, then they would not match at **e***_u_*_3_, and vice versa. To overcome this problem, it is necessary to think more carefully about how trajectories transition between basins of attraction. This issue can be dealt with rigorously (Graham and Tél, 1986; Roy and Nauman, 1995), but the general principles are outlined here. Let Ω_1_ be the basin of attraction corresponding to **e***_s_*_1_ and Ω_2_ be the basin of attraction corresponding to **e***_s_*_2_. Let ∂Ω be the separatrix between these two basins (i.e., the *x*-axis). The most probable way for a trajectory to transition from Ω_1_ to Ω_2_ involves passing through the lowest point on the surface specified by Φ_1_ along ∂Ω. Examination of Φ_1_ indicates that this point is **e***_u_*_2_. In the small-noise limit, the transition rate from Ω_1_ to Ω_2_ will correspond to Φ_1_ (**e***_u_*_2_). Similarly, the transition rate from Ω_2_ to Ω_1_ will correspond to Φ_2_ (**e***_u_*_3_). The transition rate into Ω_1_ must equal the transition rate out of Ω_2_. Therefore, the two local quasi-potentials should be translated so that the minimum heights along the separatrix are the same. In other words, one must define translated local quasi-potentials 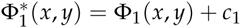 and 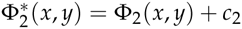 so that

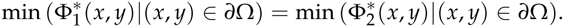

In example 1, the minima of both local quasi-potentials occurred at the same point, so the algorithm amounted to matching at that point. In example 3, the minimum saddle for Φ_1_ is **e***_u_*_2_ and the minimum saddle for Φ_2_ is **e***_u_*_3_; the heights of the surfaces at these respective points should be matched. Thus, *c*_1_ = Φ_2_(*e_u_*_3_) − Φ_1_(*e_u_*_2_) and *c*_2_ = Φ_1_(*e_u_*_2_) − Φ_2_(*e_u_*_2_). Conveniently in example 3, this is satisfied without requiring any translation (one can use *c*_1_ = *c*_2_ = 0). Finally, the global quasi-potential is found by taking the minimum value of the matched local quasi-potentials at each point. This process is automated in **QPot**, but users can also manipulate the local quasi-potential matrices manually to verify the results. This is recommended when dealing with unusual or complicated separatrices. The code below applies the automated global quasi-potential calculation to example 3.

~~~
ex3.global <- QPGlobal(local.surfaces = list(eq1.local, eq2.local),unstable.eq.x = c(0, -1.5, 1.5), unstable.eq.y = c(0, 0, 0), x.bound = bounds.x, y.bound = bounds.y)
~~~

### Step 5: Global quasi-potential visualization

Figure 14 shows a contour plot of the global quasi-potential. Note that the surface is continuous, but not smooth. The lack of smoothness is a generic feature of global quasi-potentials created from pasting local quasi-potentials. Cusps usually form when switching from the part of solution obtained from one local quasi-potential to the other.

**Figure 14:**
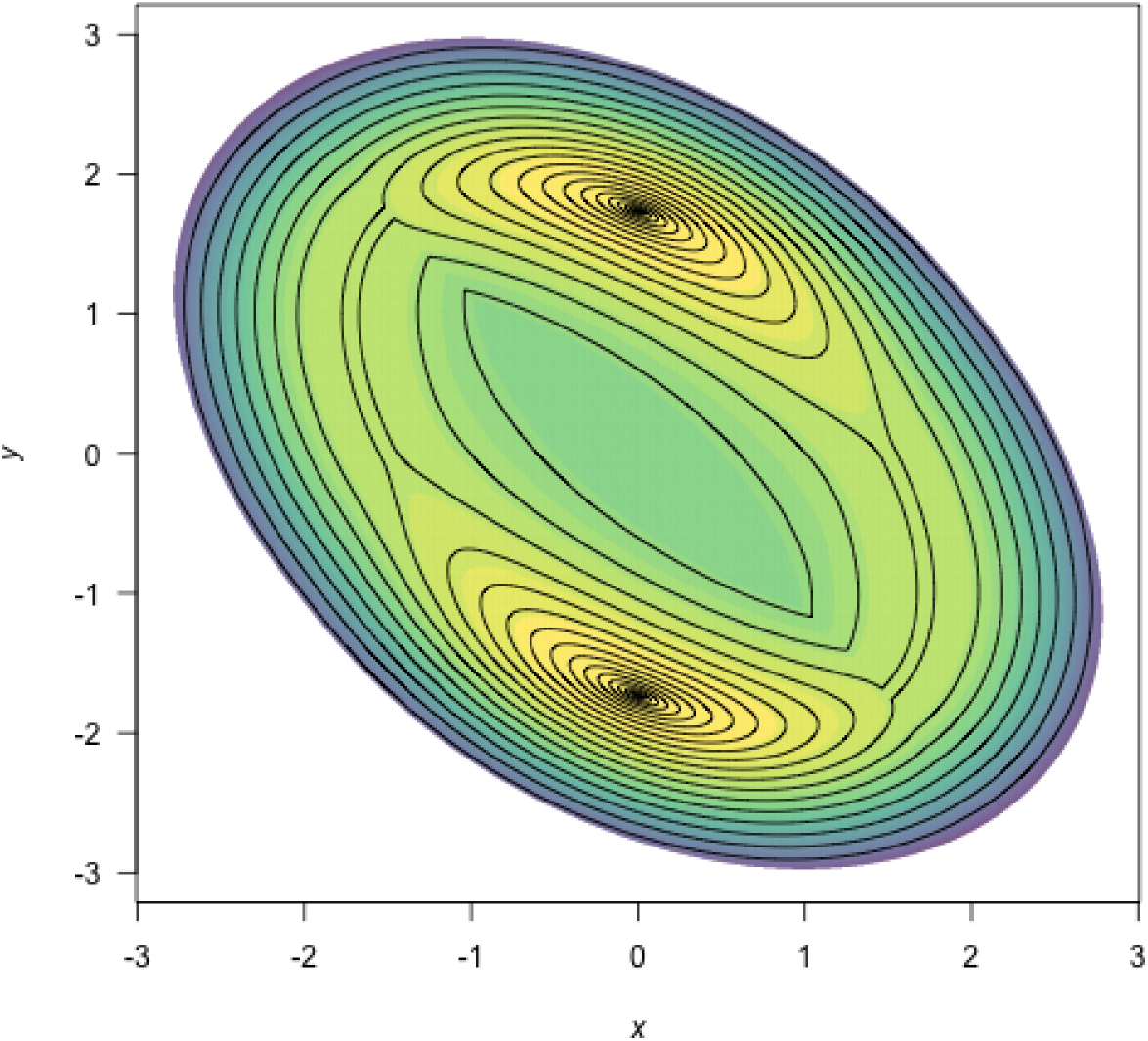
A contour plot of the quasi-potential of system (5) using the function `QPContour()`. Yellow corresponds to low values of the quasi-potential, and purple to high values.

~~~
QPContour(ex3.global, dens = c(1000, 1000), x.bound = bounds.x, y.bound = bounds.y, c.parm = 5)
~~~

## Boundary behavior

It is important to consider the type of behavior that should be enforced at the boundaries and on coordinate axes (*x* = 0 and *y* = 0). By default, the ordered-upwind method computes the quasipotential for the system defined by the user, without regard for the influence of the boundaries or the significance of these axes. In some cases, however, a model is only valid in a subregion of phase space. For example, in many population models, only the non-negative phase space is physically meaningful. In such cases, it is undesirable to allow the ordered-upwind method to consider trajectories that pass through negative phase space. In the default mode for `QPotential()`, if (*x, y*) lies in positive phase space, Φ(*x, y*) can be impacted by the vector field in negative phase space, if the path corresponding to the minimum “work” passes through negative phase space. The argument bounce = ‘d’ corresponds to this (d)efault behavior. A user can prevent the ordered upwind method from passing trajectories through negative phase space by using the option bounce = ‘p’ for (p)ositive values only. This option can be interpreted as a reflecting boundary condition. It forces the front of solutions obtained by the ordered upwind method to stay in the defined boundaries, in this case the positive phase space. A more generic option is bounce = ‘b’ for (b)ounce, which reflects based on the user-supplied boundaries. Even using this option, it is still wise to have padding space along coordinate boundaries to prevent premature termination of the algorithm, which is set with bounce.edge.

## Different noise terms

In the cases considered so far, the noise terms for the *X* and *Y* variables have had identical intensity. This was useful for purposes of illustration in the algorithm, but it will often be untrue of real-world systems. Fortunately, **QPot** can accomodate other noise terms with coordinate transforms. Consider a system of the form:

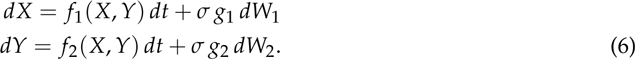

*σ* is a scaling parameter that specifies the overall noise intensity. The parameters *g*_1_ and *g*_2_ specify the relative intensity of the two noise terms. To transform this system into a form that is useable for **QPot**, make the change of variable 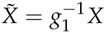 and 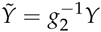. In the new coordinates, the drift terms (that is, the terms multiplied by *dt*), will be different. These new drift terms can be incorporated into the deterministic skeleton that is input into **QPot**. After obtaining the global quasi-potential for these transformed coordinates, one can switch back to the original coordinates for plotting.

Many models contain multiplicative noise terms. These are of the form:

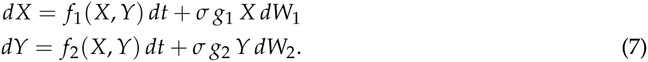

To transform this system into a form that is useable for **QPot**, make the change of variable 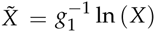 and 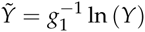. This coordinate change is non-linear, so Itô’s lemma introduces extra terms into the drift of the transformed equations. If *σ* is small, though, these terms can be discounted, and the new drift terms will remain independent of *σ*. These new drift terms can be input into **QPot**. After obtaining the global quasi-potential for these transformed coordinates, one can switch back to the original coordinates.

## Conclusion

**QPot** is an R package that provides several important tools for analyzing two-dimensional systems of stochastic differential equations. These include functions for generating realizations of the stochastic differential equations, and for analyzing and visualizing the results. A central component of **QPot** is the calculation of quasi-potential functions, which are highly useful for studying stochastic dynamics. For example, quasi-potential functions can be used to compare the stability of different attractors in stochastic systems, a task that traditional linear stability analysis is poorly suited for (Nolting and Abbott, Accepted). By offering an intuitive way to quantify attractor stability, quasi-potentials are poised to become an important means of understanding phenomena like metastability and alternative stable states. **QPot** makes quasi-potentials accessible to R users interested in applying this new framework.

## Author contributions

K.C.A, C.M.M., B.C.N., and C.R.S. designed the project. M.K.C. wrote the C code for finding the quasi-potential; C.M.M. and C.R.S. wrote the R code and adapted the C code.

## Acknowledgements

This work was supported by a Complex Systems Scholar grant to K.C.A. from the James S. McDonnell Foundation. M.K.C. was partially supported by NSF grant 1217118.

